# The synthetic triterpenoids CDDO-TFEA and CDDO-Me, but not CDDO, are potent BACH1 inhibitors

**DOI:** 10.1101/2021.10.29.466411

**Authors:** Laura Casares, Rita Moreno, Maureen Higgins, Sharadha Dayalan Naidu, Graham Neill, Lena Cassin, Anders E. Kiib, Esben B. Svenningsen, Tadashi Honda, Thomas B. Poulsen, Albena T. Dinkova-Kostova, David Olagnier, Laureano de la Vega

## Abstract

The transcription factor BACH1 is a potential target against a variety of chronic conditions linked to oxidative stress and inflammation, and formation of cancer metastasis. However, only a few BACH1 degraders/inhibitors have been described. BACH1 is a transcriptional repressor of heme oxygenase 1 (HMOX1), which is positively regulated by transcription factor NRF2 and is highly inducible by derivatives of the synthetic oleanane triterpenoid 2- cyano-3,12-dioxooleana-1,9(11)-dien-28-oic acid (CDDO). Most of the therapeutic activities of these compounds are due to their anti-inflammatory and antioxidant properties, which are widely attributed to their ability to activate NRF2. However, with such a broad range of action, these drugs may have other molecular targets that have not been fully identified and could also be of importance for their therapeutic profile. Herein we identified BACH1 as a target of CDDO-derivatives, but not CDDO. While both CDDO and CDDO-derivatives activate NRF2 similarly, only CDDO-derivatives inhibit BACH1, which explains the much higher potency of CDDO-derivatives as HMOX1 inducers compared with unmodified CDDO. Notably, we demonstrate that CDDO-derivatives inhibit BACH1 via a novel mechanism that reduces BACH1 nuclear levels while accumulating its cytoplasmic form. Altogether, our study identifies CDDO-derivatives as dual KEAP1/BACH1 inhibitors, providing a rationale for further therapeutic uses of these drugs.

## Background

The synthetic oleanane triterpenoid 2-cyano-3,12-dioxooleana-1,9(11)-dien-28-oic acid (CDDO) and its derivatives, including CDDO-methyl ester (CDDO-Me, also known as Bardoxolone methyl) and CDDO-trifluoroethyl amide (CDDO-TFEA), are a class of multifunctional drugs with anti-inflammatory and antioxidant properties that have a wide range of therapeutic uses, from neuroprotection to anticancer, in a variety of preclinical models [1–5]. These compounds were first identified as inducers of heme oxygenase 1 (HMOX1), an inducible enzyme with potent antioxidant and anti-inflammatory properties, and later as potent activators of the transcription factor NRF2 [6]. Extensive structure-activity studies led to the development of the most potent NRF2 activators known to date, some of which, including CDDO-Me, are currently in advanced clinical trials [7, 8]. NRF2 is largely controlled at the protein stability level, and its main regulator, KEAP1 (Kelch-like ECH-associated protein 1), is a substrate adaptor for the Cul3-based E3 ubiquitin ligase, and in normal conditions, KEAP1 targets NRF2 for proteasomal degradation, keeping the levels of NRF2 low in cells [9]. KEAP1 is also a sensor for electrophiles, such as CDDO and its derivatives, which chemically modify cysteines in KEAP1 [10, 11] preventing it from targeting NRF2 for degradation, leading to a rapid nuclear accumulation of NRF2 and transcription of its target genes [9].

In addition to NRF2, the transcription of *HMOX1* is also regulated by BACH1 (broad complex, tramtrack and bric à brac and cap’n’collar homology 1), a transcription factor that competes with NRF2 for binding to sequences called antioxidant response elements (AREs) within its promoter region. Unlike NRF2 which activates *HMOX1* transcription, BACH1 represses it [12–15]. While KEAP1 inhibitors/NRF2 activators induce the expression of numerous cytoprotective genes, BACH1 inhibitors/degraders activate only a limited subset of these genes, although they are extremely potent at inducing *HMOX1*.

Despite their therapeutic potential [16–23], only a few BACH1 inhibitors/degraders have been identified so far. The most widely used BACH1 degrader is hemin, a heme derivative. Hemin binds to BACH1, promoting its nuclear export and subsequent cytoplasmic degradation [24–26]. Other degraders/inhibitors are the natural phytocannabinoid cannabidiol [27], the synthetic compound HPP-4382 [19], and its derivatives [16], although their mechanisms of action are not clear. Based on the differential effect of BACH1 versus KEAP1 inhibitors, we expect drugs with dual activity, targeting both transcription factors, to have broader and stronger anti-inflammatory and antioxidant properties with potentially greater therapeutic value than drugs targeting either protein individually. In that regard, we have recently reported a chemical derivative of cannabidiol with dual activity [28], although its efficacy *in vivo* has not been established.

CDDO-derivatives are more potent than CDDO at inducing HMOX1 [6, 29] and have a better therapeutic profile, although the reason for this increased activity is unclear. In this work we demonstrate that CDDO derivatives (particularly CDDO-Me and CDDO-TFEA) are potent BACH1 inhibitors, while CDDO is not. This dual KEAP1 and BACH1 inhibition explains their enhanced potency as *HMOX1* inducers and may also explain some of their superior therapeutic profile.

## Materials and Methods

### Cell culture

Cells were grown in RPMI (HaCaT and HK2) or DMEM (H1299, A549) containing 10% FBS at 37 °C and 5% CO2. LX2 cells were maintained in high glucose DMEM media with 2mM L-Glutamine, without sodium pyruvate and with 2% FBS EmbryoMax^TM^ (Sigma-Aldrich, St. Louis, MO, USA). HaCaT cells have been validated by STR profiling and were routinely tested for mycoplasma. LX2 cells were obtained from SIGMA, and HK2, H1299 and A549 cells were obtained from ATCC and also tested for mycoplasma. CRISPR-edited NRF2-KO HaCaT cells were produced by transfecting HaCaT cells with pLentiCRISPR-v2 (a gift from Dr Feng Zhang, Addgene plasmid #52961) containing a guide RNA directed against the exon 2 of the NFE2L2 locus (which encodes NRF2) (5 -TGGAGGCAAGATATAGATCT-3 ). HaCaT BACH1-KO and HaCaT NRF2-KO/BACH1-KO cells were generated by transfecting either HaCaT WT or HaCaT NRF2-KO cells with two different pLentiCRISPR-v2 plasmids containing each one a guide RNA against the first exon and the second exon of BACH1, respectively (5 - CGATGTCACCATCTTTGTGG-3 , 5 -GACTCTGAGACGGACACCGA-3 ). All CRISPR-edited cell lines were selected with puromycin for 2 days, cells were clonally selected by serial dilution, and positive clones were identified as previously described [30]. Control cells, referred as HaCaT wild type (HaCaT WT), are the pooled population of surviving cells transfected with an empty pLentiCRISPRv2 vector treated with puromycin.

### Antibodies and reagents

Antibodies against BETA-ACTIN (C-4), BACH1 (F-9) and LAMIN B2 (C-20) were obtained from Santa Cruz Biotechnology (Dallas, Texas, USA). Anti-NRF2 (D1Z9C) was obtained from Cell Signalling Technology (Danvers, MA, USA) and anti-HMOX1 was purchased from Biovision (San Francisco, CA, USA). Antibody against ALPHA-TUBULIN was obtained from Sigma- Aldrich (St. Louis, MO, USA). HRP-conjugated secondary antibodies were obtained from Life Technologies (Carlsbad, California, USA). Dimethyl sulfoxide (DMSO) was from Sigma- Aldrich. R,S-sulforaphane (SFN) was purchased from LKT Laboratories (St. Paul, MN, USA). (±)-TBE-31 was synthesized as described [31, 32]. CDDO and CDDO-derivatives were obtained from Cayman Chemicals (Ann Arbor, MI, USA). MG132 was obtained from Santa Cruz Biotechnology; Leptomycin B from Cayman Chemicals, MLN4924 and Selinexor (KPT-330) from Selleckchem (Houston, TX, USA) and Actinomycin D and Cycloheximide from Sigma.

### Plasmids

BACH1-RFP, and BACH1- C435, C46, C492, C646A (Hemin resistant) -RFP were generated as follows. BACH1 WT or Hemin-resistant inserts were synthesised and cloned into Plenti-CMV-MCS-RFP-SV-puro. Plenti-CMV-MCS-RFP-SV-puro was a gift from Jonathan Garlick & Behzad Gerami-Naini (Addgene plasmid # 109377; http://n2t.net/addgene:109377; RRID:Addgene_109377).

### Quantitative real time PCR (rt-qPCR)

RNA from cells was extracted using GeneJET RNA Purification Kit (Thermo Fisher Scientific) and 500 ng of RNA per sample was reverse-transcribed to cDNA using Omniscript RT kit (Qiagen) supplemented with RNase inhibitor according to the manufacturer’s instructions. Resulting cDNA was analysed using TaqMan Universal Master Mix II (Life Technologies, Carlsbad, CA, USA) as well as corresponding Taqman probes. Gene expression was determined using a QuantStudio 7 Flex qPCR machine by the comparative ΔΔCT method. All experiments were performed at least in triplicates and data were normalized to the housekeeping gene HPRT1. Taqman probes used: HPRT1 Hs02800695_m1; HMOX1 Hs01110250_m1; AKR1B10 Hs00252524_m1.

### Cell lysis and western blot

Cells were washed and harvested in ice-cold phosphate-buffered saline (PBS). For <u>whole cell extracts,</u> cells were lysed in RIPA buffer supplemented with phosphate and protease inhibitors [50 mM Tris- HCl pH 7.5, 150 mM NaCl, 2 mM EDTA, 1% NP40, 0.5% sodium deoxycholate, 0.5 mM Na3VO4, 50 mM NaF, 2 μg/mL leupeptine, 2 μg/mL aprotinin, 0.05 mM pefabloc]. Lysates were sonicated for 15 s at 20% amplitude and then cleared by centrifugation for 15 min at 4 °C. For <u>subcellular fractionation,</u> cells were resuspended in 400 μl of low-salt buffer A (10 mM Hepes/KOH pH7.9, 10 mM KCL, 0.1 mM EDTA, 0.1 mM EGTA, 1 mM β-Mercaptoethanol) and after incubation for 10 min on ice, 10 μl of 10% NP-40 was added and cells were lysed by gently vortexing. The homogenate was centrifuged for 10 s at 13,200 rpm, the supernatant representing the cytoplasmic fraction was collected and the pellet containing the cell nuclei was washed 4 additional times in buffer A. The pellet containing the nuclear fraction was then resuspended in 100 μl high-salt buffer B (20 mM Hepes/KOH pH7.9, 400 mM NaCL, 1 mM EDTA, 1 mM EGTA, 1 mM β-mercaptoethanol). The lysates were sonicated and centrifuged at 4 °C for 15 min at 13,200 rpm. The supernatant representing the nuclear fraction was collected. Protein concentration was determined using the BCA assay (Thermo Fisher Scientific, Waltham, MA, USA). Lysates were mixed with SDS sample buffer and boiled for 7 min at 95 °C. Equal amounts of protein were separated by SDS-PAGE, followed by semidry blotting to a polyvinylidene difluoride membrane (Thermo Fisher Scientific). After blocking of the membrane with 5% (w/v) non-fat dried milk dissolved in Tris buffered saline (TBS) with 0.1% v/v Tween-20 (TBST), membranes were incubated with the primary antibodies overnight at 4°C. Appropriate secondary antibodies coupled to horseradish peroxidase were detected by enhanced chemiluminescence using ClarityTM Western ECL Blotting Substrate (Bio-Rad, Hercules, CA, USA). Resulting protein bands were quantified and normalised to each lane’s loading control using the ImageStudio Lite software (LI-COR). For whole cell extracts, the protein of interest was normalised against ACTIN or GADPH. LAMIN was used as an internal control for nuclear extracts and TUBULIN or GADPH were used as controls for cytoplasmic extracts.

### Cell viability assay

Alamar Blue (Thermo Fisher Scientific) was used to determine cell viability after drug treatment. HaCaT cells were seeded in 96-well plates to 50–60% confluency and treated the next day with the corresponding compounds for 48 hours. After treatment, Alamar Blue was added to the wells (1:10 ratio) and after four hours of incubation at 37 °C the fluorescence was measured (excitation 550 and an emission at 590 nm) using a microplate reader (Spectramax m2). Viability was calculated relative to the DMSO treated control.

### Statistical analysis

Experiments were repeated at least 2-5 times with multiple technical replicates to be eligible for the indicated statistical analyses. Data were analysed using Graphpad Prism statistical package. All results are presented as mean ± SD unless otherwise mentioned. The differences between groups were analysed using one-way ANOVA.

## Results

### CDDO-derivatives, but not CDDO, reduce BACH1 levels

We have previously shown in immortalised human keratinocytes (HaCaT cells), that the classical NRF2 activator sulforaphane (SFN) is a weak *HMOX1* inducer (but a very good inducer of the NRF2 transcriptional target *AKR1B10*), while BACH1 degraders such as hemin strongly induce *HMOX1* (in an NRF2-independent manner) without affecting *AKR1B10* expression [27, 28]. This emphasizes that although *HMOX1* has often been used as a surrogate for NRF2 activity, in some cases *AKR1B10* induction might be a more appropriate reporter for NRF2 activation while *HMOX1* induction is a better surrogate for BACH1 inhibition. To answer whether the observed limited effect of SFN on *HMOX1* in HaCaT cells is a general phenomenon for NRF2 activators we compared three potent NRF2 activators (SFN, CDDO and TBE31) against a BACH1 degrader (hemin) for their ability to induce *HMOX1* in these cells. As shown in figure 1A, all three NRF2 activators were weak *HMOX1* inducers when compared with hemin but potent inducers of *AKR1B10* expression.

**Figure 1.**
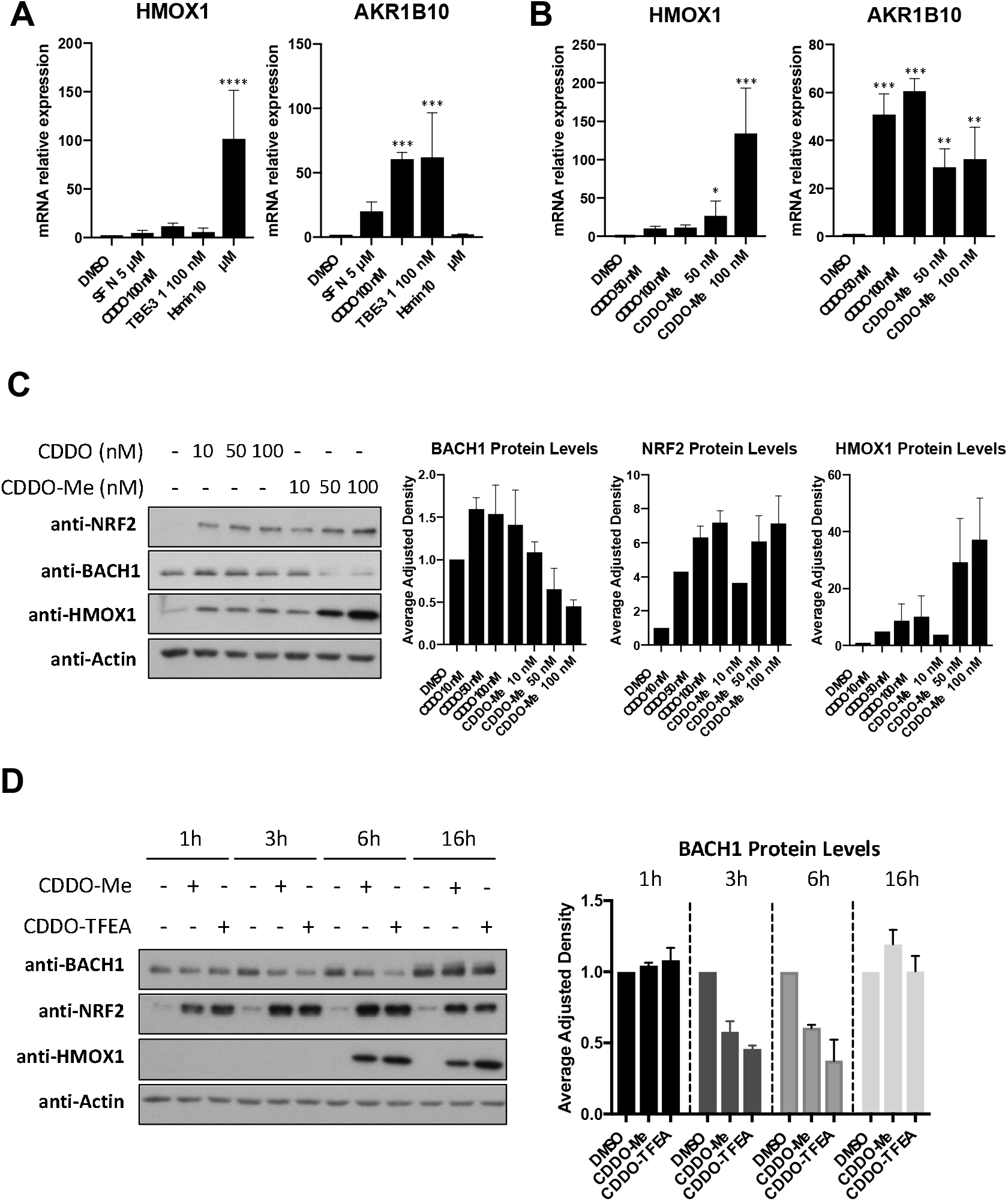
CDDO derivatives, but not CDDO, reduce BACH1 levels. **(A)** HaCaT cells were treated with either DMSO (0.1%, v/v), SFN (5 μM), CDDO (100 nM), TBE-31 (100 nM) or Hemin (10 μM) for 16h. Cells were lysed and mRNA levels of *HMOX1* and *AKR1B10* were analysed by qRT-PCR, using *HPRT1* as a housekeeping gene. **(B)** As in A, but HaCaT cells were treated with either DMSO (0.1%, v/v) or increasing concentrations of CDDO or CDDO-Me. After 16 hours cells were harvested and lysed and mRNA levels of *HMOX1* and *AKR1B10* were analysed by real-time qPCR. Data were normalised using *HPRT1* as an internal control (n= 3) and are expressed relative to the DMSO treated sample. **(C)** HaCaT cells were treated with DMSO (0.1%, v/v) or increasing concentrations of CDDO or CDDO-Me. Five hours later, cells were harvested and lysed. Protein levels of NRF2, BACH1, HMOX1 and ACTIN were analysed by Western Blot. Left panel shows a representative blot and right panels show quantification of NRF2 and HMOX1 protein levels against the loading control. Data represent means ± SD (n= 3) and are expressed relative to the DMSO-treated samples. **(D)** HaCaT cells were treated with either DMSO (0.1%, v/v), CDDO-Me (100 nM) or CDDO-TFEA (100 nM) for 1h, 3h, 6h or 16h. Cells were harvested, lysed and analysed for the levels of the indicated proteins. Left panel is a representative blot; right panels are the quantification of BACH1 levels (n= 3). Data are expressed relative to the DMSO-treated samples for each time point and were normalized against their respective loading controls (DMSO sample for each time point set to 1). *P ≤ 0.05, **P ≤ 0.01, ***P ≤ 0.001, ****P ≤ 0.0001.

Since CDDO-Me is more potent than CDDO at inducing *HMOX1* expression in some cellular models, we tested whether CDDO-Me and CDDO had a differential effect on *HMOX1* transcription in HaCaT cells. CDDO-Me was significantly more potent than CDDO at inducing *HMOX1* expression, although both compounds were equally potent at inducing *AKR1B10* (Fig. 1B), suggesting that their differential effect on HMOX1 must be NRF2-independent. Next, we hypothesised that, in addition to activating NRF2, CDDO-Me might be targeting BACH1. To test this, we compared the effect that CDDO and CDDO-Me had on BACH1 and NRF2 protein levels. As shown in figure 1C, CDDO-Me - but not CDDO - reduced BACH1 protein levels and greatly induced HMOX1, while both compounds equally stabilised NRF2. Since other CDDO-derivatives are also potent *HMOX1* inducers, we hypothesised that they might also reduce BACH1 protein levels. To test this, we compared the effect of various CDDO-derivatives on BACH1 and NRF2 protein levels as well as *HMOX1* and *AKR1B10* expression. Of the derivatives tested, CDDO-TFEA and CDDO-Me were the most potent compounds at reducing BACH1 levels (Suppl. Fig S1A) and at inducing *HMOX1* expression (Suppl. Fig S1B). All compounds (CDDO and derivatives) induced *AKR1B10* to a similar extent (Suppl. Fig S1B). Based on their potency, we focused on CDDO-TFEA and CDDO-Me (structures shown in Suppl. Fig S1C) and performed a time course analysis of their effect on BACH1 levels. Our results show that BACH1 reduction appears to be maximal between three and six hours, and that this effect is not observed at 16 hours (Fig. 1 D). Neither CDDO-TFEA nor CDDO-Me reduced cell viability at the concentrations used in various cellular systems (Suppl. Fig. S1D).

### The differential effect of CDDO and CDDO-derivatives on *HMOX1* expression is due to BACH1 inhibition

Reportedly, some CDDO-derivatives still increase HMOX1 protein levels in the absence of NRF2 [29], although the factor responsible for that induction has not been identified. To test whether in our system the differential effect of CDDO-TFEA and CDDO-Me versus CDDO was dependent on NRF2, we compared wild type (WT) and NRF2-KO HaCaT cells. We found that both CDDO-TFEA and CDDO-Me were more potent than CDDO at inducing *HMOX1* in WT cells, and a similar pattern (although with reduced fold induction) was observed in NRF2-KO cells (Fig. 2A), demonstrating that the differential effect between CDDO and CDDO-TFEA/Me was indeed not related to NRF2. On the other hand, *AKR1B10* induction in WT cells was similar for the three compounds and was completely abolished in the absence of NRF2 (Fig. 2B). We used a complementary approach with an immortalised human proximal tubular kidney cell line (HK2) to test if the observed NRF2-independent differential effect of CDDO-derivatives on *HMOX1* was cell-type specific. Using CRISPR/Cas9 gene editing, we produced an isogenic HK2 cell line with hyperactive NRF2 that cannot be further stabilised by activators (NRF2-GOF cells) (Cell line validation in Suppl. Fig S2A). In this cell line, CDDO failed to induce *HMOX1* any further while CDDO-Me and CDDO-TFEA still potently induced *HMOX1* (Suppl. Fig. S2B), confirming that this differential induction does not depend on NRF2 stabilisation. In agreement with the results obtained in HaCaT cells, the three compounds equally induced *AKR1B10* in WT HK2 cells but failed to induce it further in NRF2-GOF HK2 cells (Suppl. Fig. S2C).

**Figure 2.**
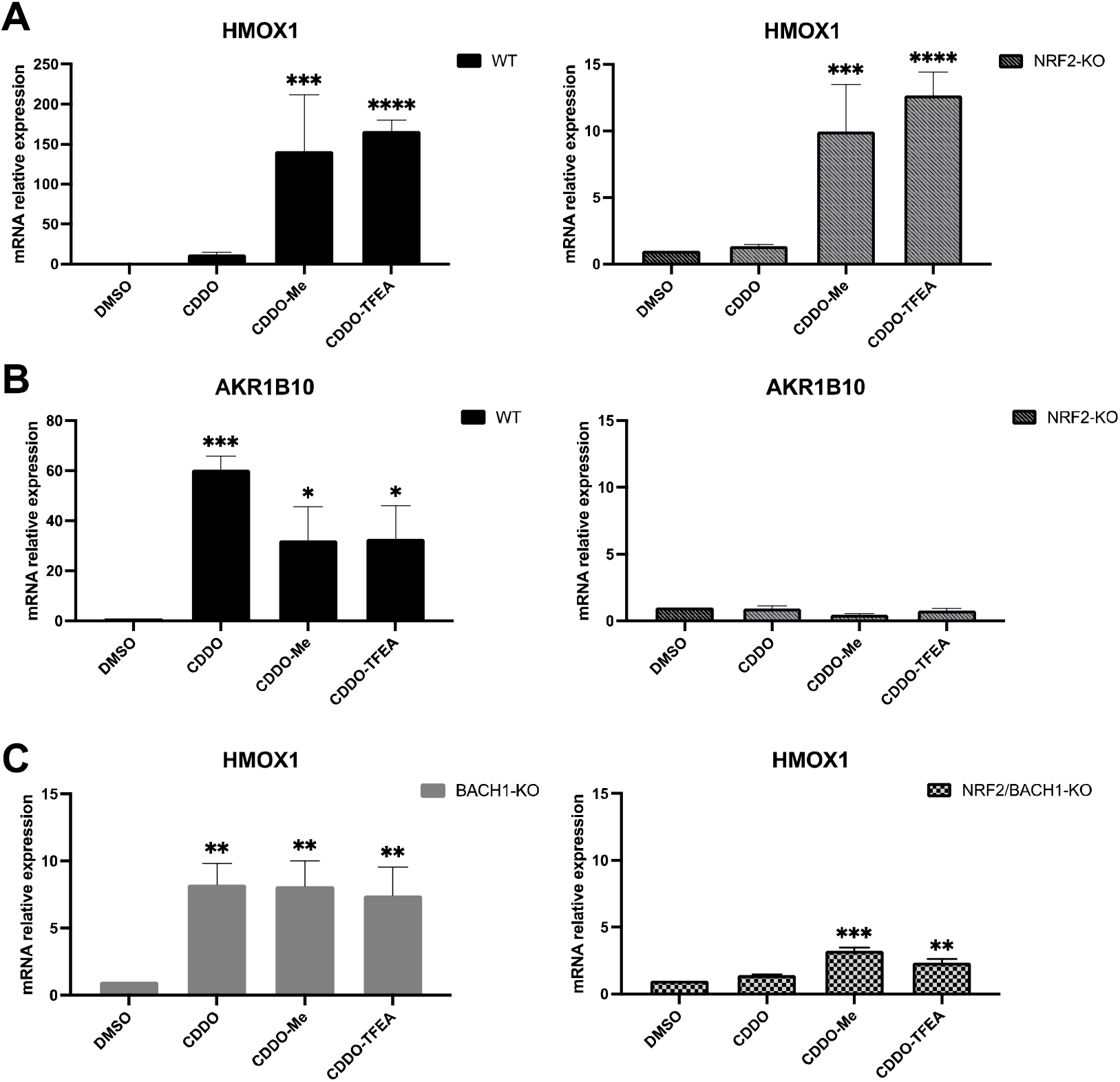
The differential effect of CDDO and CDDO/TFEA on *HMOX1* is BACH1-dependent and NRF2-independent. **(A,B)** HaCaT WT or NRF2-KO cells were treated with either DMSO (0.1%, v/v), CDDO (100 nM), CDDO-Me (100 nM) or CDDO-TFEA (100 nM) for 16h. Samples were collected and mRNA levels of *HMOX1* (A) and *AKR1B10* (B) were analysed via real-time qPCR, using *HPRT1* as an internal control. Data are expressed relative to the DMSO-treated samples in each cell line (DMSO in WT and NRF2-KO cells set to 1). **(C)** HaCaT BACH1-KO and HaCaT NRF2/BACH1-KO cells were treated as above. Levels of *HMOX1* were analysed by qRT-PCR as previously described. *HMOX1* levels in the DMSO samples of each cell line were set to 1 and the rest of the data are expressed relative to their corresponding DMSO sample. *P ≤ 0.05, **P ≤ 0.01, ***P ≤ 0.001, ****P ≤ 0.0001.

As BACH1 is a key regulator of *HMOX1* expression, we hypothesised that the differential effect between CDDO and CDDO-derivatives must be due to their differential activity on BACH1, and that the strong effect of CDDO-derivatives on *HMOX1* expression relates to the combination of NRF2 stabilisation and BACH1 reduction. To test this, we compared the three compounds in BACH1-KO and in BACH1/NRF2-KO HaCaT cells. In BACH1-KO cells, the differential effect between CDDO and CDDO-derivatives on *HMOX1* was lost (Fig. 2C **left panel)**, suggesting that BACH1 is indeed responsible for that effect (comparison between Fig. 2C and 2A), and that NRF2 (or another factor) might be responsible for the remaining observed induction. In fact, in double BACH1/NRF2-KO cells the effect of the compounds on *HMOX1* expression was largely abolished, highlighting the relevance of both factors regulating *HMOX1* (Fig. 2C **right panel**).

### CDDO-derivatives reduce BACH1 nuclear levels while accumulating cytoplasmic BACH1 levels in a NRF2-independent manner

Our results demonstrate that CDDO-derivatives, but not CDDO, reduce the levels of BACH1, and that this reduction is responsible for their differential effect on *HMOX1* expression. However, the reduction of BACH1 levels was less than expected based on the strong *HMOX1* induction (which was similar to that observed with the potent BACH1 degrader hemin). Furthermore, although CDDO-derivatives were still robust inducers of *HMOX1* in other cell lines (such as HK2 cells, the lung cancer cells H1299 and A549, or the human hepatic stellate cell line LX2), in contrast to HaCaT cells (Fig. 1B), they did not affect the BACH1 levels (Suppl. Fig. S3A-S3D), which was intriguing. As some of the compounds that target BACH1 for degradation do so by first inducing its nuclear export [25], we wondered whether CDDO-derivatives might affect the balance between nuclear/cytoplasmic BACH1 and whether the compound effect could be on nuclear BACH1 (the active pool). Indeed, while in HK2, LX2, H1299 and A549 cells the effect of CDDO- derivatives on total BACH1 (whole cell extract) was insignificant (Suppl. Fig. S3A-S3D), by using subcellular fractionation we observed that CDDO-Me and CDDO-TFEA significantly reduced nuclear BACH1 while increasing its cytoplasmic levels (Fig. 3A-3D). This explains the apparent lack of effect on total BACH1 levels (as the cytoplasmic accumulation would mask its nuclear reduction) and the strong *HMOX1* induction (as nuclear BACH1 represents the transcriptionally active pool).

**Figure 3.**
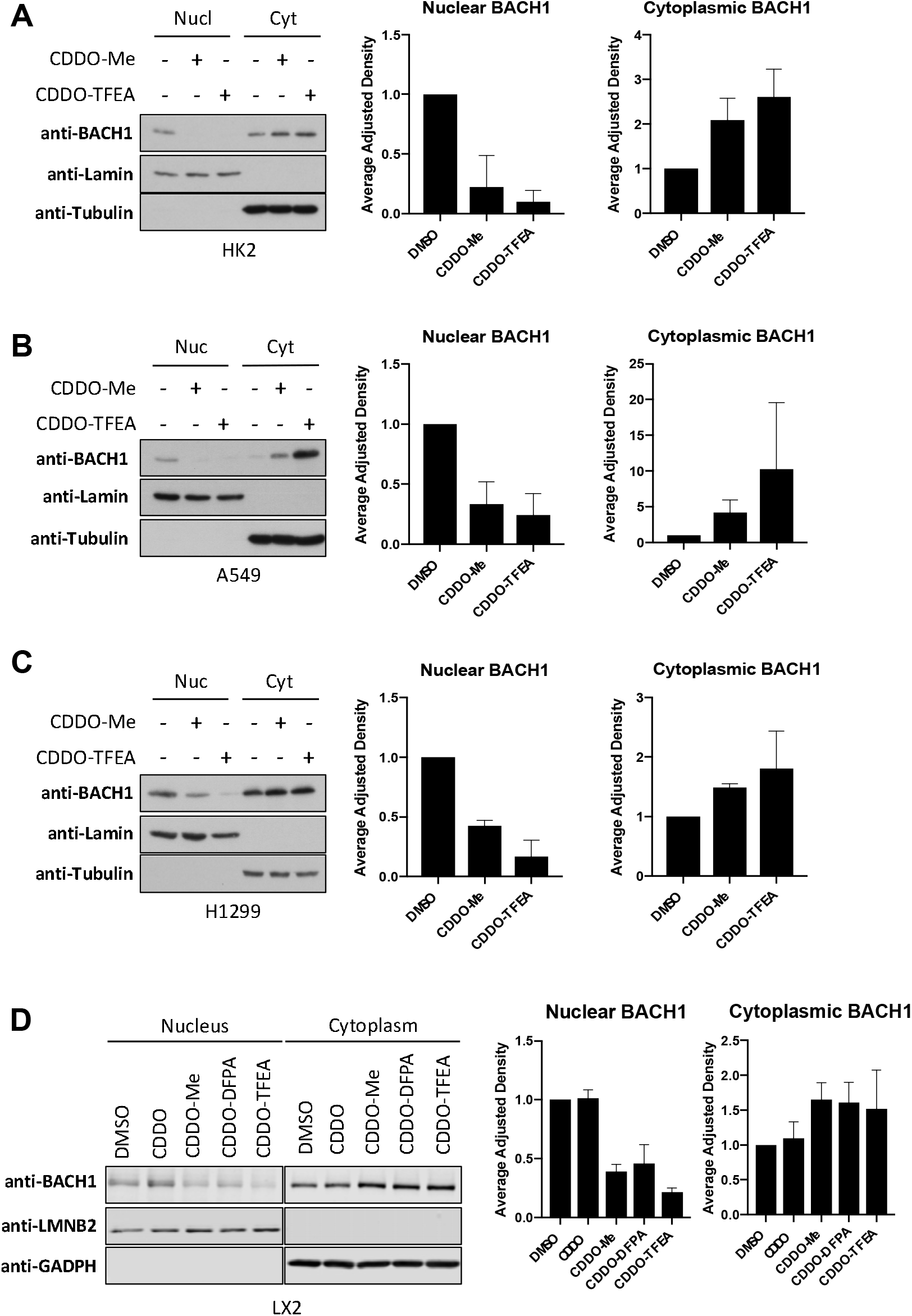
CDDO-derivatives reduce nuclear BACH1 levels while increasing cytoplasmic BACH1 levels. **(A-D)** HK2 (A), A549 (B), H1299 (C) or LX2 (D) cells were treated with DMSO (0.1%, v/v), CDDO-Me (100 nM) or CDDO-TFEA (100 nM). Six hours later cells were harvested and subcellular fractionation was performed. BACH1 protein levels were analysed via western blot. Panels on the left show a representative blot; panels on the right are the corresponding BACH1 nuclear and cytoplasmic quantifications, which were normalised against their internal control (i.e., LAMIN for nuclear and TUBULIN for cytoplasmic levels). Data represent means ± SD (n= 3) and are expressed relative to the DMSO-treated samples.

Additionally, as these compounds are potent NRF2 activators and NRF2 induces BACH1 expression [33, 34], we tested whether NRF2 was necessary for the effect of CDDO- derivatives on BACH1 nuclear and cytoplasmic levels. To do this, we performed time course experiments in WT and NRF2-KO HaCaT cells. The absence of NRF2 did not impair the reduction in nuclear BACH1 nor its cytoplasmic accumulation (comparison between Suppl Fig. S3E and S3F), strongly suggesting that NRF2 is not required for either of these effects. In agreement, potent NRF2 activators such as CDDO or TBE31 did not induce BACH1 cytoplasmic accumulation or promote its nuclear reduction (Suppl Fig. S3G).

### How are CDDO-derivatives affecting BACH1 levels?

The nuclear reduction and cytoplasmic accumulation of BACH1 in response to CDDO-TFEA/Me could be explained in different ways:

1- The two effects are not linked: e.g. CDDO-derivatives induce BACH1 nuclear degradation and independently, BACH1 cytoplasmic accumulation, either by increasing the protein stability or the transcript levels of BACH1.
2- The two effects are linked: e.g. CDDO-derivatives affect the nuclear export of BACH1.

We next tested these two possible explanations:

### Is BACH1 protein stability affected by the CDDO-derivatives?

The two main pathways controlling protein degradation are the ubiquitin-proteasome system and autophagy. To study the involvement of the ubiquitin-proteasome system we used MG132 (proteasome inhibitor) and MLN 4924 (an inhibitor of NEDD8 activating enzyme, which acts by inhibiting all Cullin RING ligases). Although both inhibitors increased the basal levels of BACH1, neither of them abolished the effect of CDDO-TFEA/Me on BACH1 (Fig. 4A), suggesting that degradation of BACH1 via the proteasome is not the main mechanism by which the CDDO- derivatives reduce levels of BACH1. To address the potential role of autophagy, we used the autophagy inhibitor bafilomycin A1, which did not impair the effect of CDDO-Me/TFEA on BACH1 protein levels (Suppl Fig. S4A).

**Figure 4.**
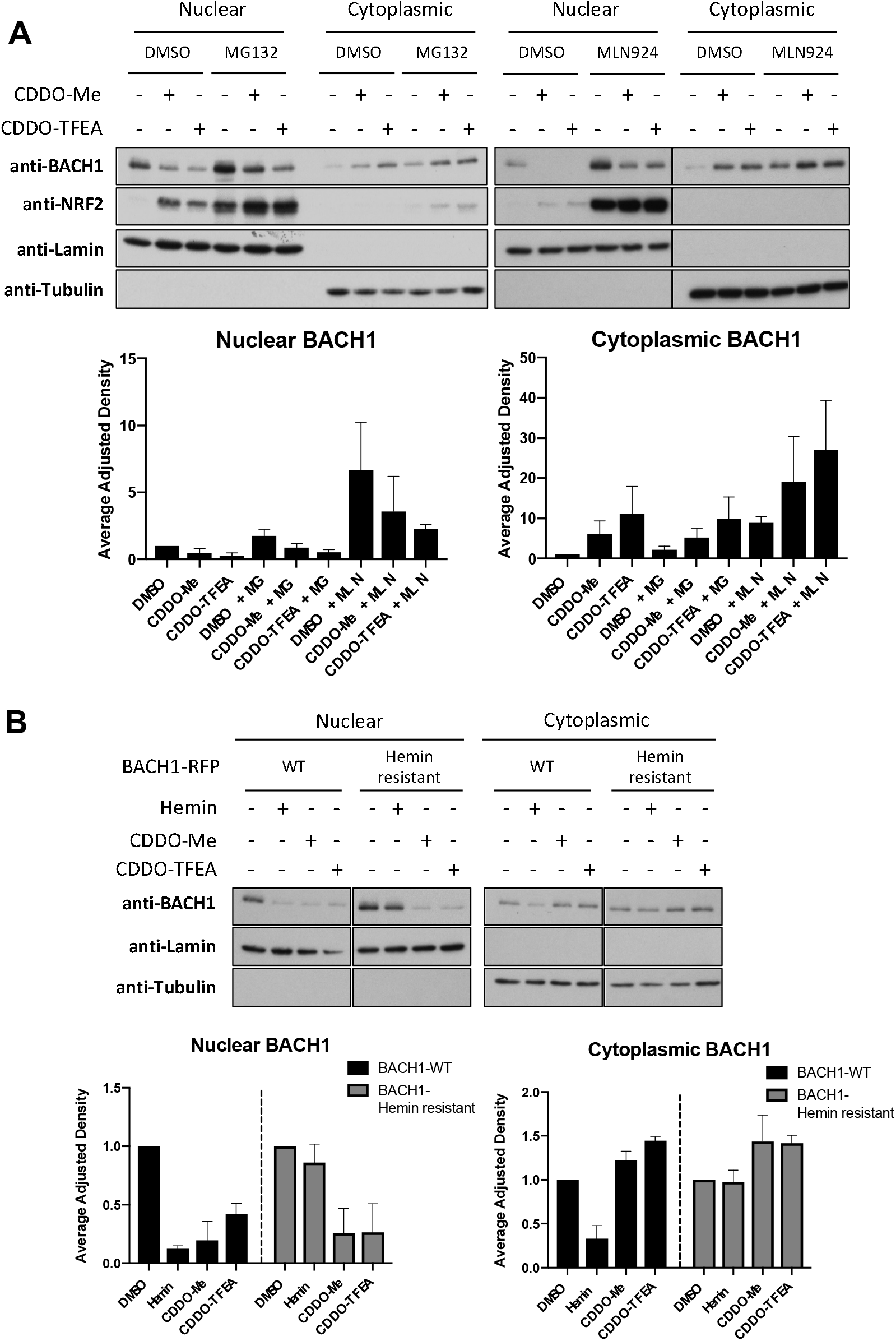
CDDO-Me/TFEA affects BACH1 levels in a proteasome independent manner and have a mechanism of action different than hemin. **(A)** HaCaT cells were incubated with either DMSO (0.1%, v/v), MG132 (10 μM) or MLN924 (2 μM) for one hour. After that, either DMSO (-), CDDO-Me (100 nM) or CDDO-TFEA (100 nM) was added. Six hours later, cells were harvested and nuclear/cytoplasmic fractions were isolated and analysed for their levels of BACH1 and NRF2. Upper panel is a representative blot and lower panels are the quantifications of nuclear and cytoplasmic BACH1 levels normalised against their corresponding loading control. Data represent means ± SD (n= 3) and are expressed relative to the DMSO sample. **(B)** HaCaT BACH1-KO cells reconstituted with either BACH1- RFP-WT or BACH1-RFP-Hemin resistant mutant were treated with DMSO (-), Hemin (10 μM), CDDO-Me (100 nM) or CDDO-TFEA (100 nM) for six hours. Cells were harvested and nuclear/cytoplasmic fractions were isolated and analysed for their levels of BACH1. Upper panel is a representative blot and lower panels are the quantifications of nuclear and cytoplasmic BACH1 levels normalised against their corresponding loading control. Data represent means ± SD (n= 3) and are expressed relative to their DMSO control.

Hemin (the best-characterised BACH1 degrader) binds to BACH1, promoting its proteasomal degradation, and thus our results suggest that CDDO-derivatives and hemin might have different mechanisms of action. To address this, we reconstituted BACH1-KO cells with either BACH1-WT or a BACH1 hemin-resistant mutant, in which four cysteines in the haem-binding site were mutated to alanine (Hemin-resistant) [25, 35]. Although both hemin and the CDDO-derivatives efficiently reduced nuclear levels of BACH1-WT, only CDDO-TFEA/Me reduced the levels of the hemin-resistant BACH1 mutant (Fig. 4B). These results further confirm that the mechanism of BACH1 reduction by CDDO-TFEA/Me is different from that of hemin.

### Is BACH1 transcription affected by the CDDO-derivatives?

As BACH1 protein stability did not seem to be affected by CDDO-TFEA/Me, we tested if BACH1 transcriptional regulation was responsible for its cytoplasmic accumulation. To test this, we used compounds to inhibit either protein synthesis (cycloheximide, a protein synthesis inhibitor) or transcription (actinomycin D, a DNA-directed RNA synthesis inhibitor). Neither of these inhibitors blocked BACH1 nuclear reduction or its cytoplasmic accumulation in response to CDDO-TFEA/Me (Suppl Fig. S4B), suggesting that synthesis of new proteins (and their transcription) is not needed for the effect of the CDDO-derivatives on BACH1.

### Is BACH1 nuclear export affected by the CDDO-derivatives?

So far, our results showed that neither transcriptional regulation nor protein degradation are mechanisms responsible for the effect of CDDO-TFEA/Me, suggesting that the CDDO-derivatives might be reducing nuclear BACH1 and accumulating its cytoplasmic pool via a nuclear export mechanism. While small molecules (20-40 kD) can passively diffuse between the nucleus and the cytoplasm, transport of larger molecules such as proteins involves signal-dependent mechanisms. Many nuclear export substrates contain a nuclear export signal (NES) that binds the export receptor CRM1 (exportin 1/Xpo1), which is sensitive to inhibitors such as leptomycin B and selinexor. However, not all proteins that shuttle between the nucleus and cytoplasm use CRM1 to do so, and CRM1-independent nuclear export pathways have been identified [36–40]. To address whether the changes in nuclear and cytoplasmic BACH1 in response to CDDO-TFEA/Me are related to a CRM1-dependent nuclear export mechanism, we tested the effect of two CRM1 inhibitors (leptomycin B and selinexor) (Suppl. Fig S4C). Although both inhibitors induced a basal nuclear accumulation of BACH1, as expected, neither of them abolished its nuclear reduction nor its cytoplasmic accumulation in response to CDDO-TFEA/Me. Overall, our data suggest that CDDO-derivatives induce BACH1 nuclear export in a CRM1-independent manner.

## Discussion

Our results demonstrate that CDDO-derivatives - but not CDDO - inhibit BACH1, explaining their greater potency as *HMOX1* inducers in comparison with CDDO. Although we did not identify the mechanism(s) responsible for the ability of CDDO-derivatives to reduce nuclear BACH1 levels, our data show that neither the proteasome nor the nuclear export receptor CRM1 are involved, demonstrating that CDDO-derivatives use a mechanism different from the one used by hemin. This highlights the need to better understand the mechanisms controlling BACH1 regulation. Additionally, the differential effect observed in the levels of nuclear and cytoplasmic BACH1 shows that nuclear reduction of BACH1 (without further cytoplasmic degradation) is sufficient for a strong *HMOX1* induction. This should be taken into consideration in the design of screening strategies to identify BACH1 inhibitors, as only looking at total BACH1 levels in cells could be misleading. It would be interesting to address whether the accumulation of cytoplasmic BACH1 may have other functions that are unrelated to its well-characterized role as transcriptional regulator.

Our data demonstrate that both CDDO-TFEA and CDDO-Me are very potent dual KEAP1 and BACH1 inhibitors, which could explain some of their therapeutic benefits. Importantly our study provides a rationale for their potential clinical development for conditions affected by BACH1, such as bone destructive diseases [16], non-alcoholic steatohepatitis [22], atherosclerosis [23], insulin resistance [20], coronary artery disease [41], aging related conditions [17] and tumour metastasis [33, 34, 42–46].

## AUTHORS CONTRIBUTION

LC, RM, MH, SDN, GN, AEK, EBS, WL and LC conducted the experiments and were responsible for initial data analysis, figure preparation and statistical analysis. TH and TBP provided resources and technical expertise. LV, DO and ADK had a leading contribution in the design of the study, and an active role in the discussion and interpretation of the whole dataset. LV wrote the original draft of the manuscript. ADK, DO and LC reviewed and edited the manuscript. Funding acquisition LV, DO and ADK. All the authors take full responsibility for the work.

---

**Figure S1.**
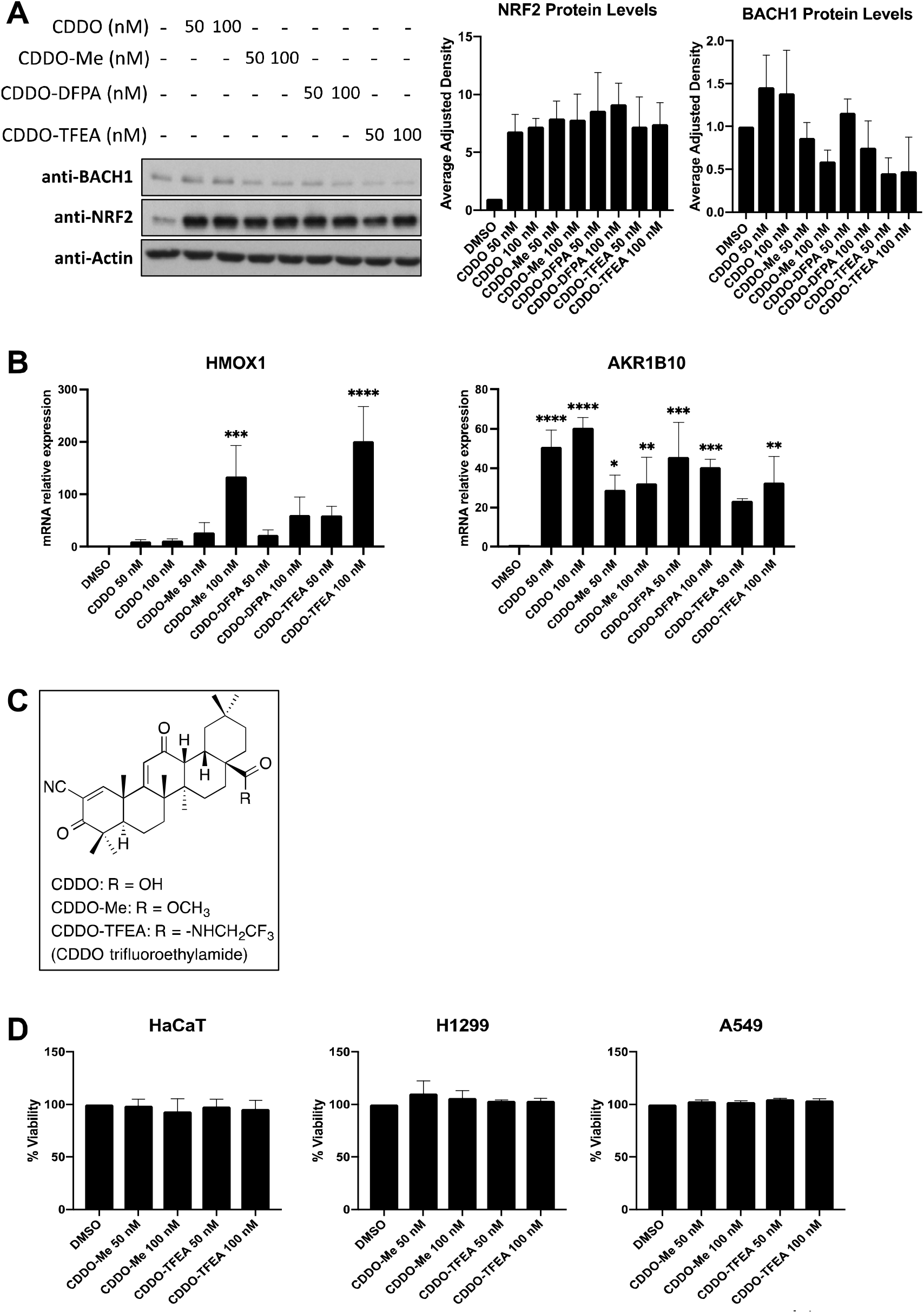
**(A)** HaCaT cells were treated with vehicle (DMSO, 0.1%, v/v) or increasing concentrations of either CDDO, CDDO-Me, CDDO-DFPA or CDDO-TFEA. After five hours cells were lysed and samples were analysed by Western Blot. Upper panel is a representative blot and lower panels show the quantification of BACH1 and NRF2 protein levels normalized for actin levels. Data represent means ± SD (n = 3) and are expressed relative to the DMSO-treated samples. **(B)** HaCaT cells were treated with either DMSO (0.1%, v/v) or different concentrations of CDDO, CDDO-Me, CDDO-DFPA or CDDO-TFEA. After 16 hours cells were harvested and lysed and mRNA levels of *HMOX1* and *AKR1B10* were analysed by real-time qPCR. Data were normalised using *HPRT1* as an internal control (n= 3) and are expressed relative to the DMSO treated sample. *P ≤ 0.05, **P ≤ 0.01, ***P ≤ 0.001, ****P ≤ 0.0001. **(C)** Structures for CDDO, CDDO-Me and CDDO-TFEA. **(D)** HaCaT, H1299 or A549 cells were treated with either DMSO (0.1%, v/v) or different concentrations of CDDO-Me or CDDO-TFEA as indicated. After 48 hours, viability was calculated relative to the DMSO-treated control using Alamar Blue.

**Figure S2.**
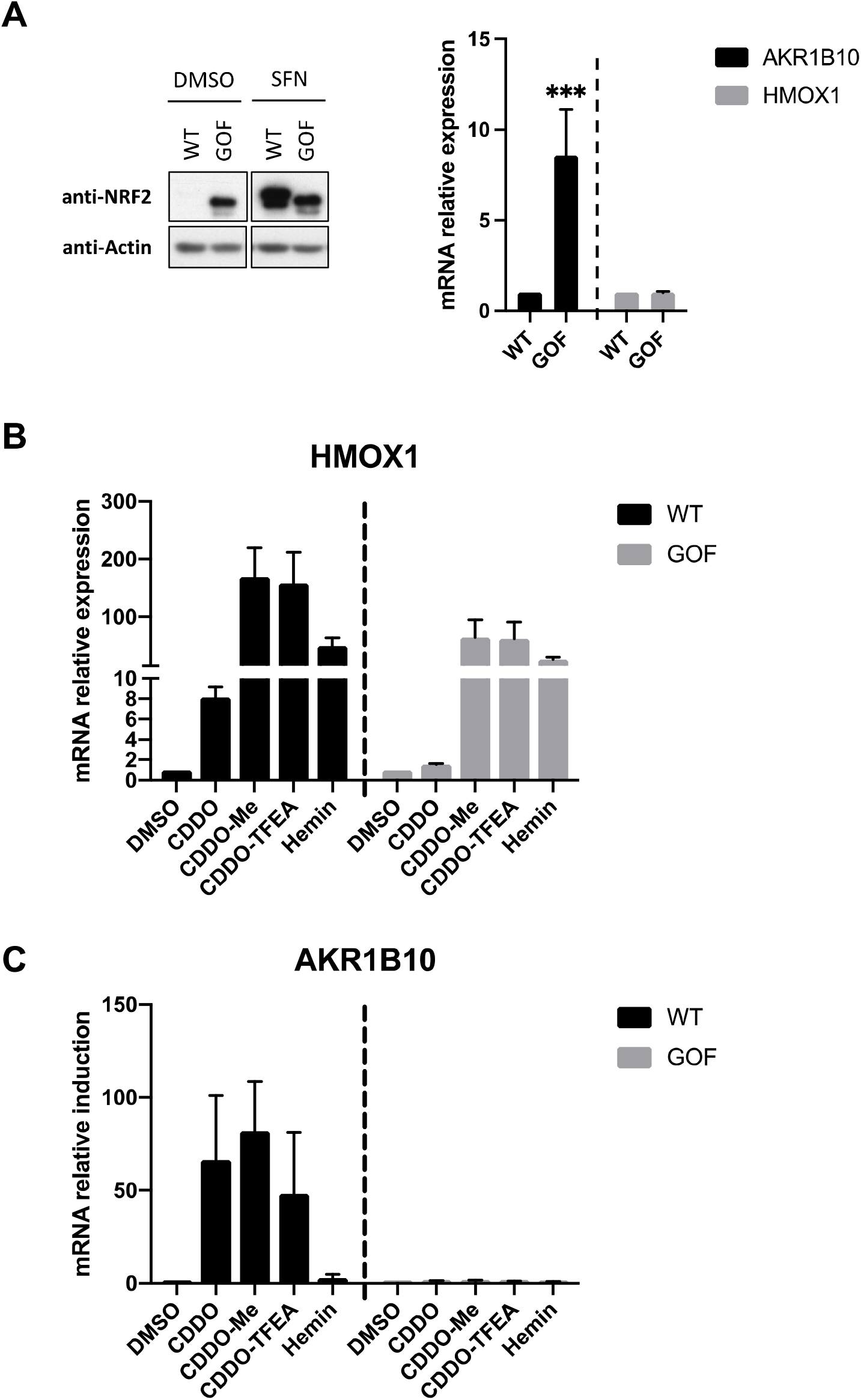
**(A)** Left panel: Control (WT) and NRF2 gain-of-function (GOF) HK2 cells were treated with either DMSO (0.1%, v/v) or sulforaphane (SFN). Three hours later the levels of NRF2 were measured by western blot. Right panel: Basal mRNA levels of *HMOX1* and *AKR1B10* in control (WT) and NRF2-GOF HK2 cells were analysed by RT-qPCR. *HPRT1* was used as a housekeeping gene for the analysis. ***P ≤ 0.001. **(B,C)** HK2 Control (WT) and NRF2-GOF cells were treated with DMSO, CDDO (100 nM), CDDO-Me (100 nM), CDDO-TFEA (100 nM) or Hemin (10 μM) for 16h. *HMOX1* (B) and *AKR1B10* (C) mRNA levels were analysed using RT-qPCR and *HPRT1* as a housekeeping gene. Data are expressed relative to the DMSO-treated samples in each cell line (DMSO in WT and NRF2-GOF cells set to 1). *P ≤ 0.05.

**Figure S3.**
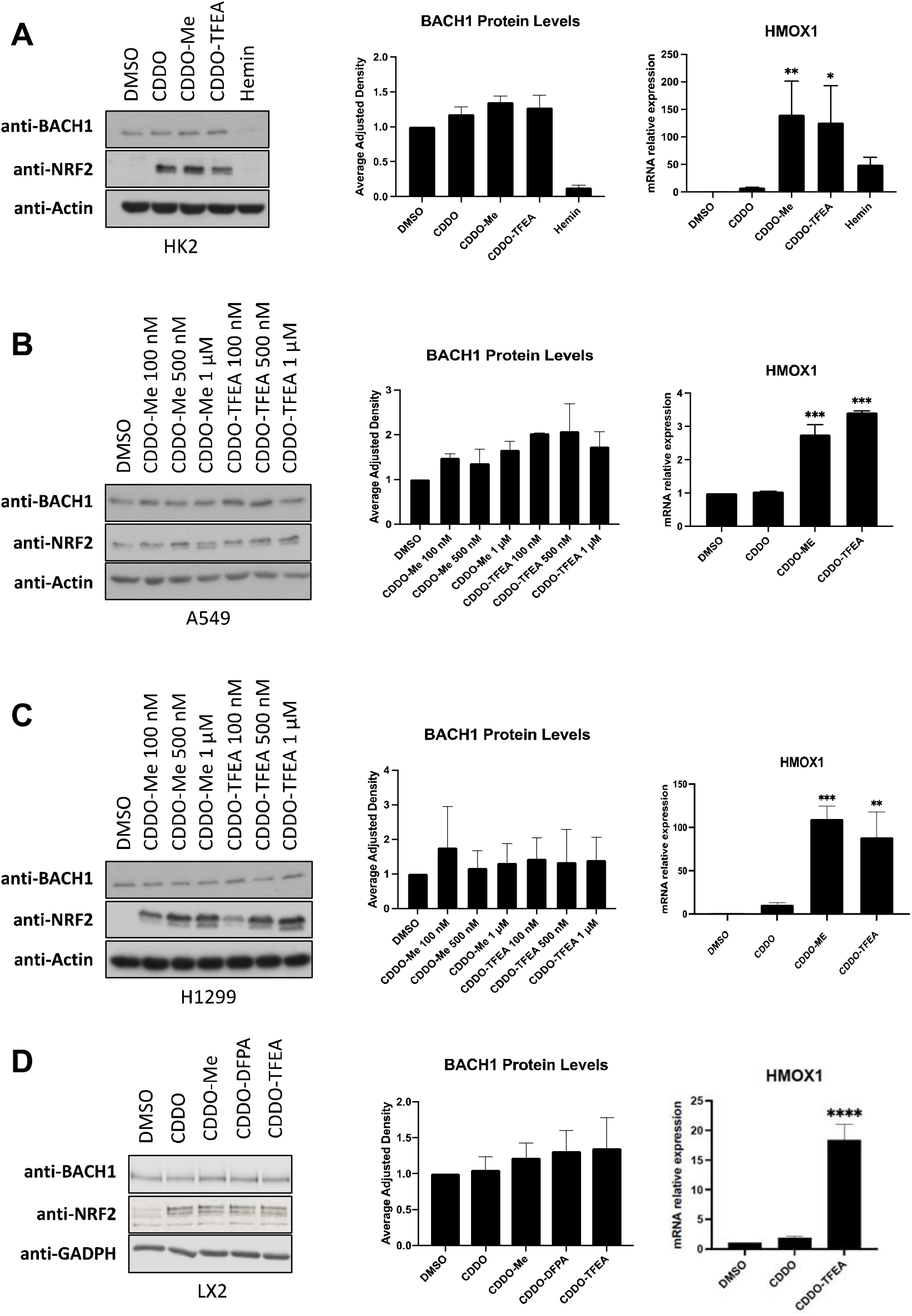
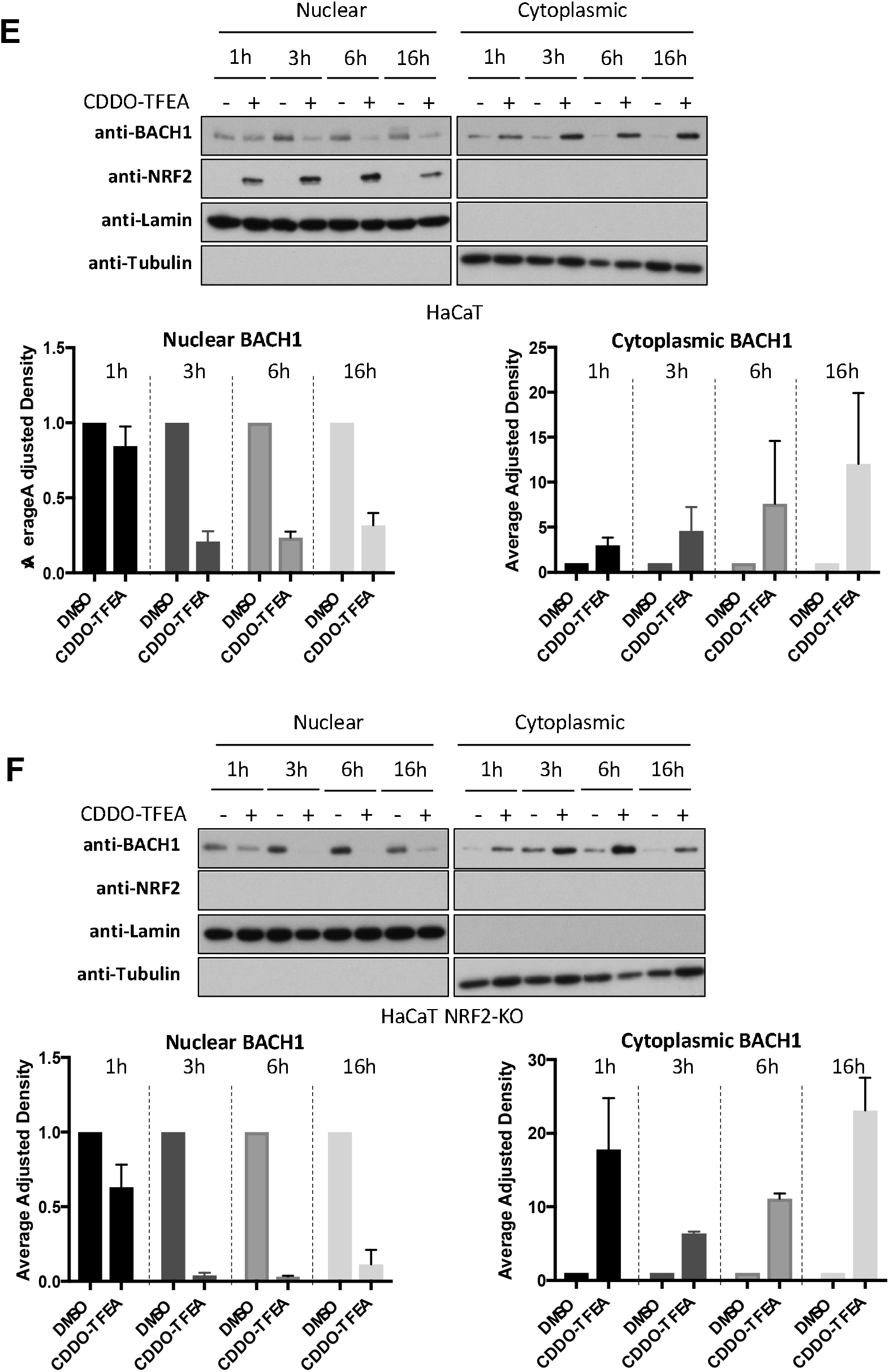
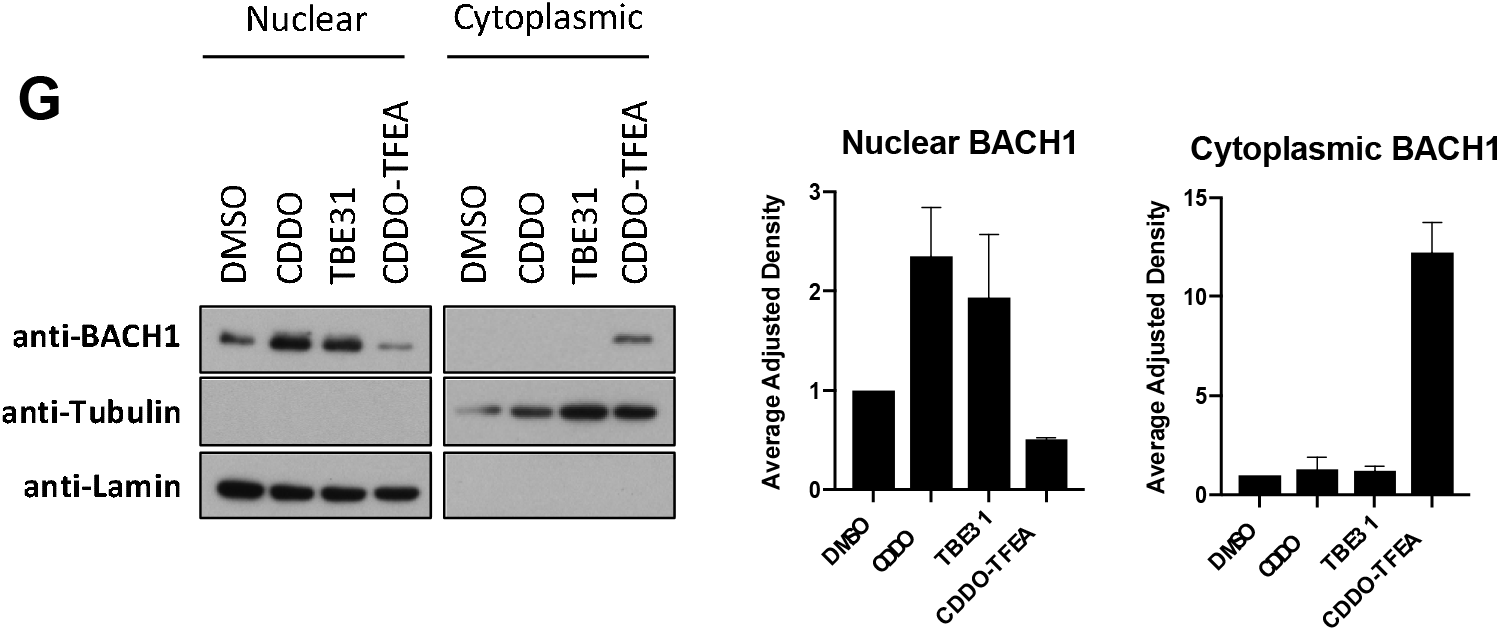
**(A)** HK2 cells were treated with DMSO, CDDO (100 nM), CDDO-Me (100 nM), CDDO-TFEA (100 nM) or hemin (10 μM) for 6h. Cells were lysed and total BACH1, NRF2 and actin levels were analysed by western blot. Representative blot is shown on the left panel and quantification of BACH1 protein levels (n=2) is in the middle panel. Right panel: HK2 cells were treated with either DMSO (0.1%, v/v), CDDO (100 nM), CDDO-Me (100 nM), CDDO-TFEA (100 nM) or hemin (10 μM) for 16h. *HMOX1* mRNA levels were analysed. *P ≤ 0.05, **P ≤ 0.01. **(B)** A549 cells were treated with DMSO (0.1%, v/v) or increasing concentrations of CDDO-Me or CDDO-TFEA for 6h. Left panel shows a representative blot while middle panels show quantification of BACH1 protein levels (n= 2). Right panel: A549 cells were treated with DMSO (0.1%, v/v), CDDO (100 nM), CDDO-Me (100 nM) or CDDO-TFEA (100 nM) for 6h. *HMOX1* mRNA levels were analysed. ***P ≤ 0.001 **(C)** H1299 cells were treated with DMSO (0.1%, v/v) or increasing concentrations of CDDO-Me or CDDO-TFEA for 16h. Left panel shows a representative blot while middle panels show quantification of BACH1 protein levels (n=2). Right panel: H1299 were treated with DMSO (0.1%, v/v), CDDO (100 nM), CDDO-Me (100 nM) or CDDO-TFEA (100 nM) for 16h. *HMOX1* mRNA levels were analysed. **P ≤ 0.01, ***P ≤ 0.001. **(D)** LX2 cells were treated with DMSO (0.1%, v/v), CDDO (50 nM), CDDO-Me (50 nM), CDDO-DFPA (50 nM) or CDDO-TFEA (50 nM) for 6h. Samples were lysed and total BACH1, NRF2 and ACTIN levels were analysed by western blot. Representative blot is shown on the left panel and quantification of BACH1 protein levels (n= 3) is in the middle panel. Right panel: LX2 cells were treated with either DMSO (0.1%, v/v), CDDO (50 nM) or CDDO-TFEA (50 nM) for 16h. *HMOX1* mRNA levels were analysed. *P ≤ 0.05, **P ≤ 0.01. **(E,F)** HaCaT WT cells (E) and NRF2-KO cells (F) were treated with DMSO (0.1%, v/v) or CDDO-TFEA (100 nM) for 1h, 3h, 6h or 16h. Cells were harvested and nuclear and cytosolic fractions were isolated and analysed for the levels of the indicated proteins. Upper panel is a representative blot; lower panels are the quantification of BACH1 nuclear and cytoplasmic levels (n= 2). Data are expressed relative to the DMSO-treated samples for each time point and were normalized against their respective loading controls (DMSO sample for each time point set to 1). **(G)** HaCaT WT cells were treated with either DMSO (0.1%, v/v), CDDO (100 nM), TBE-31 (100 nM) or CDDO-TFEA (100 nM) for six hours. Nuclear and cytosolic fractions were isolated and analysed for their levels of BACH1. Upper panel is a representative blot and lower panels show the quantification of BACH1 nuclear and cytoplasmic levels (n=2), normalized against their respective loading controls. Data is expressed relative to the DMSO treated samples.

**Figure S4.**
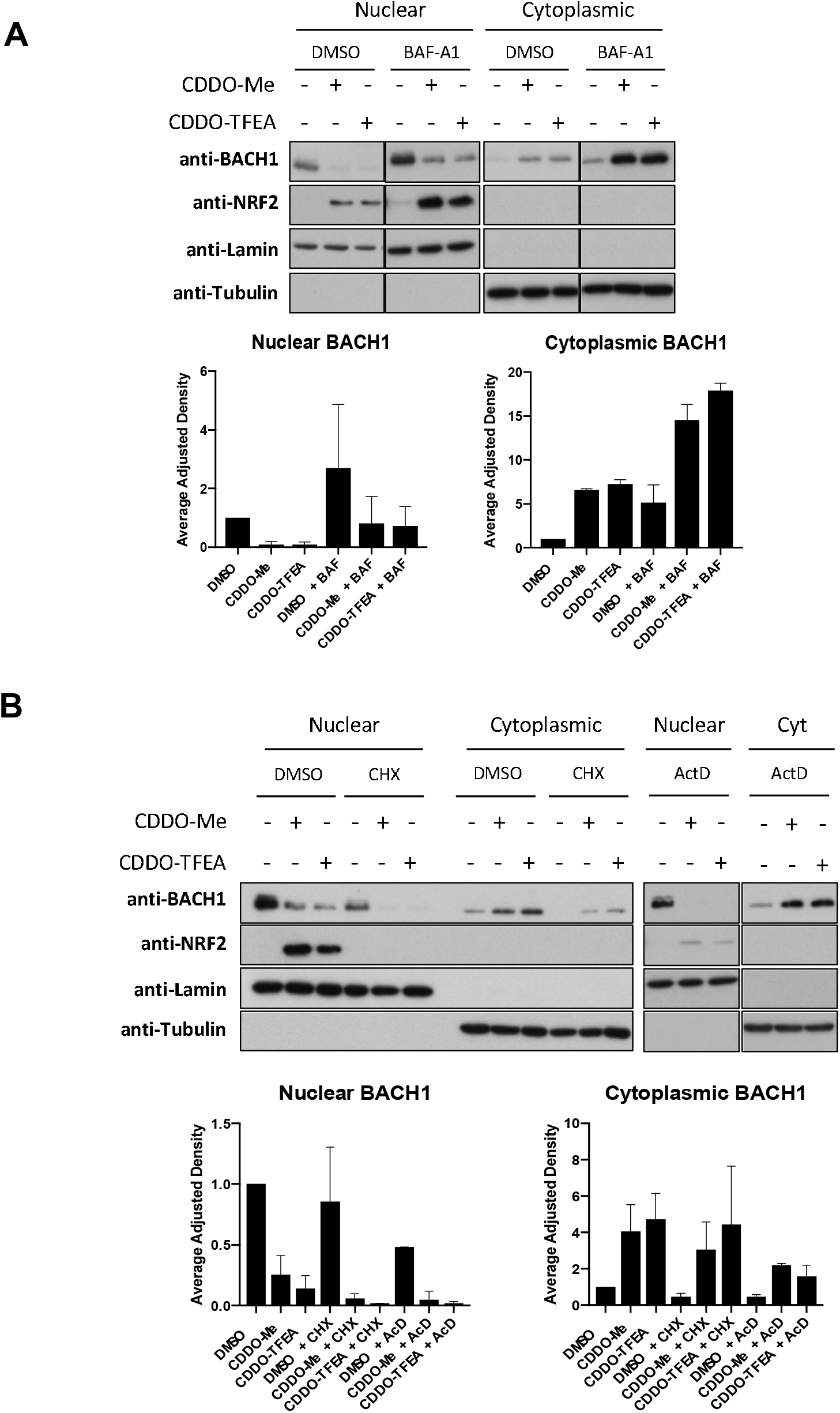
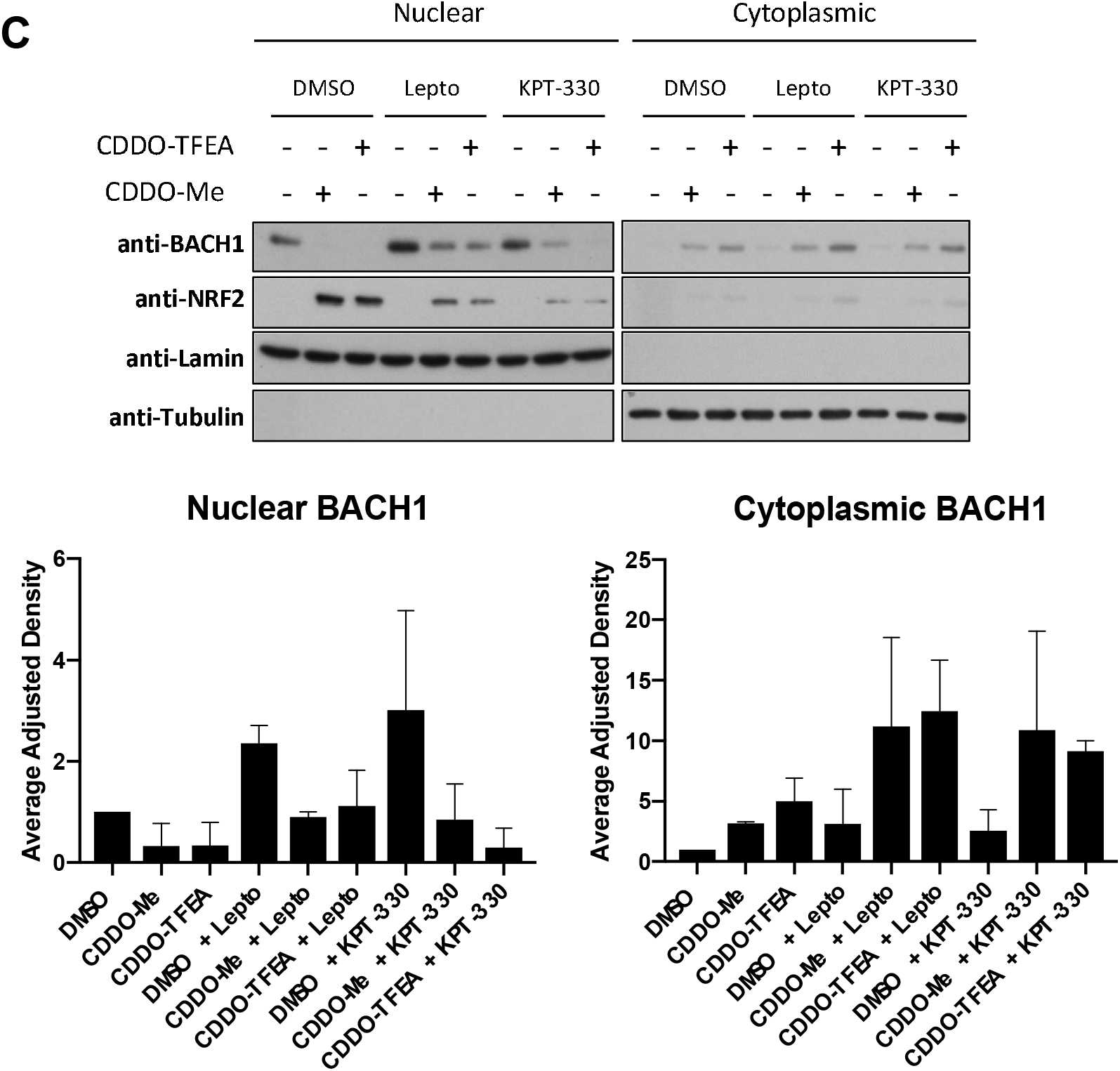
**(A)** HaCaT cells were incubated with either DMSO (0.1%, v/v) or bafilomycin A1 (BAF-A1, 100 nM). Two hours later they were treated with either DMSO (-), CDDO-Me (100 nM) or CDDO-TFEA (100 nM) for another six hours. Subcellular fractionation was performed as previously described. Upper panel is a representative blot and lower panels are the quantifications of nuclear and cytoplasmic BACH1 levels normalised against their corresponding loading control. Data represent means ± SD (n= 2) and are expressed relative to the DMSO sample. Control and BAF-A1 treated samples were all loaded in the same gel **(B)** HaCaT cells were incubated with either DMSO (0.1%, v/v), cycloheximide (CHX, 10 μM) for 2h or actinomycin D (ActD, 1 μg/mL) for 30 min. After that, either DMSO (-), CDDO-Me (100 nM) or CDDO-TFEA (100 nM) was added. Six hours later, cells were harvested and nuclear/cytoplasmic fractions were isolated and analysed for their levels of BACH1 and NRF2. Upper panel is a representative blot; lower panels are the quantifications of nuclear and cytoplasmic BACH1 levels normalised against their corresponding loading control. Data represent means ± SD (n= 3) and are expressed relative to the DMSO-treated cells. **(C)** HaCaT cells were incubated with either DMSO (0.1%, v/v), leptomycin B (Lepto, 25 ng/mL) or KPT-330 (1 μM). After two hours, either DMSO (-), CDDO-Me (100 nM) or CDDO-TFEA (100 nM) was added. Six hours later, cells were harvested and subcellular fractionation was performed. BACH1 and NRF2 protein levels were analysed by western blot. Upper panel is a representative blot and lower panels are the quantification of BACH1 nuclear and cytoplasmic levels normalised to the corresponding loading control. Data represent means ± SD (n=3) and are expressed relative to the DMSO-treated cells.

